## Supplemental Figure 1 for "Hypothalamic representation of the imminence of predator threat detected by the vomeronasal organ in mice"

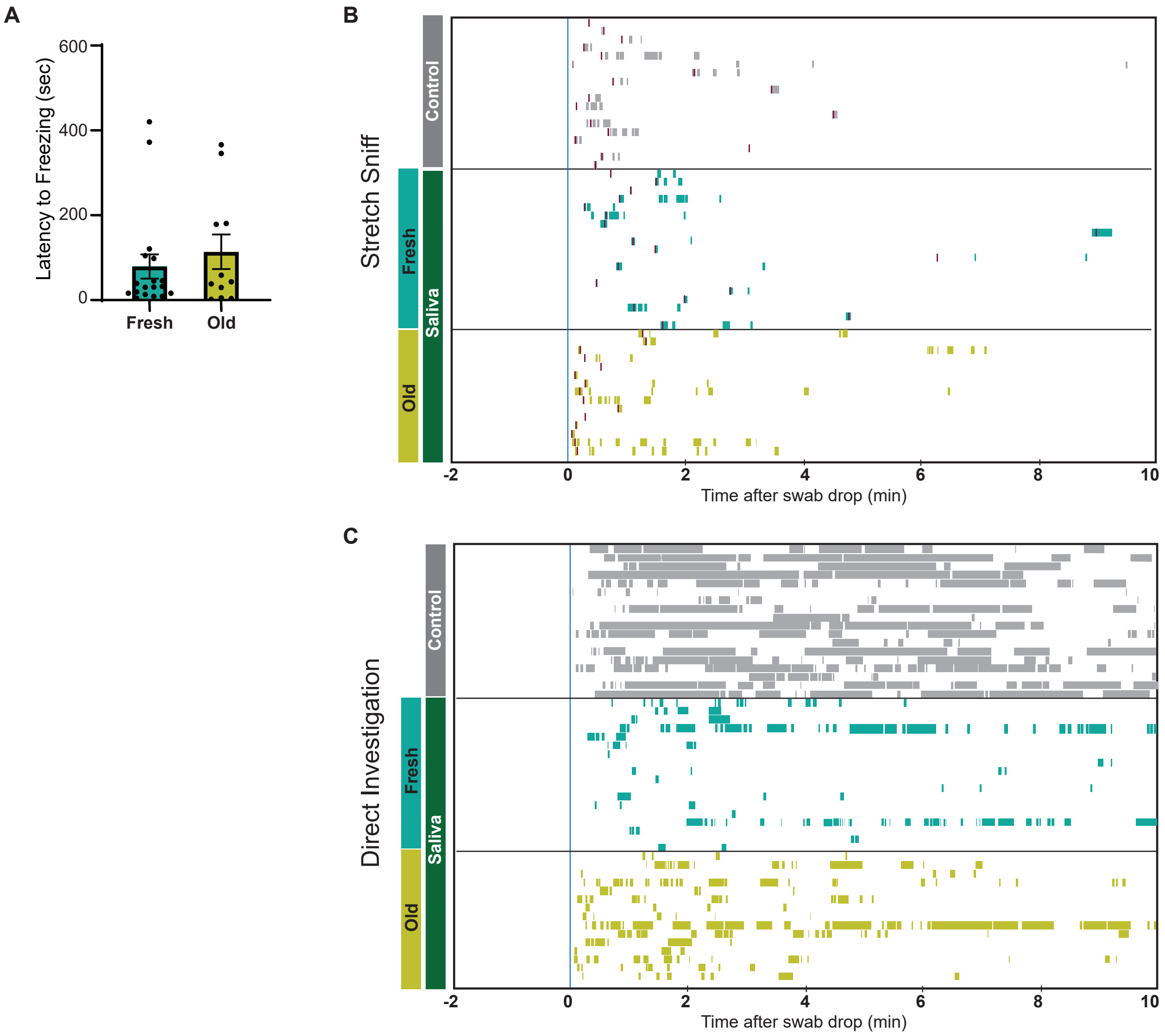


**Supplemental figure 1**

(A) Latency to the first freezing episode in mice exhibited freezing after the first interaction. Two-tailed t test (t=0.7172, df=27, p = 0.4794). (B, C) Raster plots displaying stretch sniffing (B) and direct investigation (C) episodes of individual mice exposed to a clean control swab (gray), fresh saliva (green), or old saliva (yellow). The introduction of a clean, fresh, or old saliva swab into a mouse's cage is denoted as time 0. Red lines in B indicate the first contact with a swab for each mouse.
