## Supplemental Figure 2 for "Hypothalamic representation of the imminence of predator threat detected by the vomeronasal organ in mice"

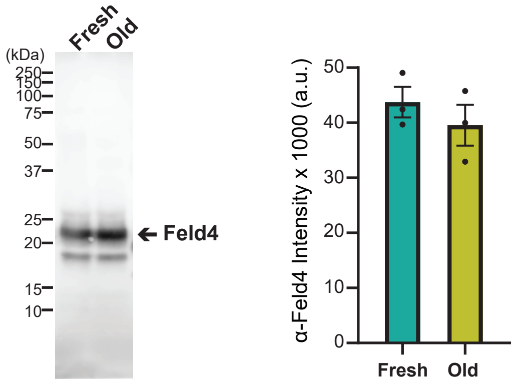


**Supplemental Figure 2**

Representative western blot analysis of Fel d 4 protein bands in Fresh and Old saliva samples. (Left) Immunoblot showing bands of Fel d 4 protein detected using anti-Fel d 4 antibody. Each lane was loaded with 10 μg of total protein. (Right) Quantification of Fel d 4 band intensities in Fresh and Old saliva samples (n = 3 each). The values are presented as means ± S.E.M., with individual dots representing individual saliva samples.
