## Supplemental Figure 3 for "Hypothalamic representation of the imminence of predator threat detected by the vomeronasal organ in mice"

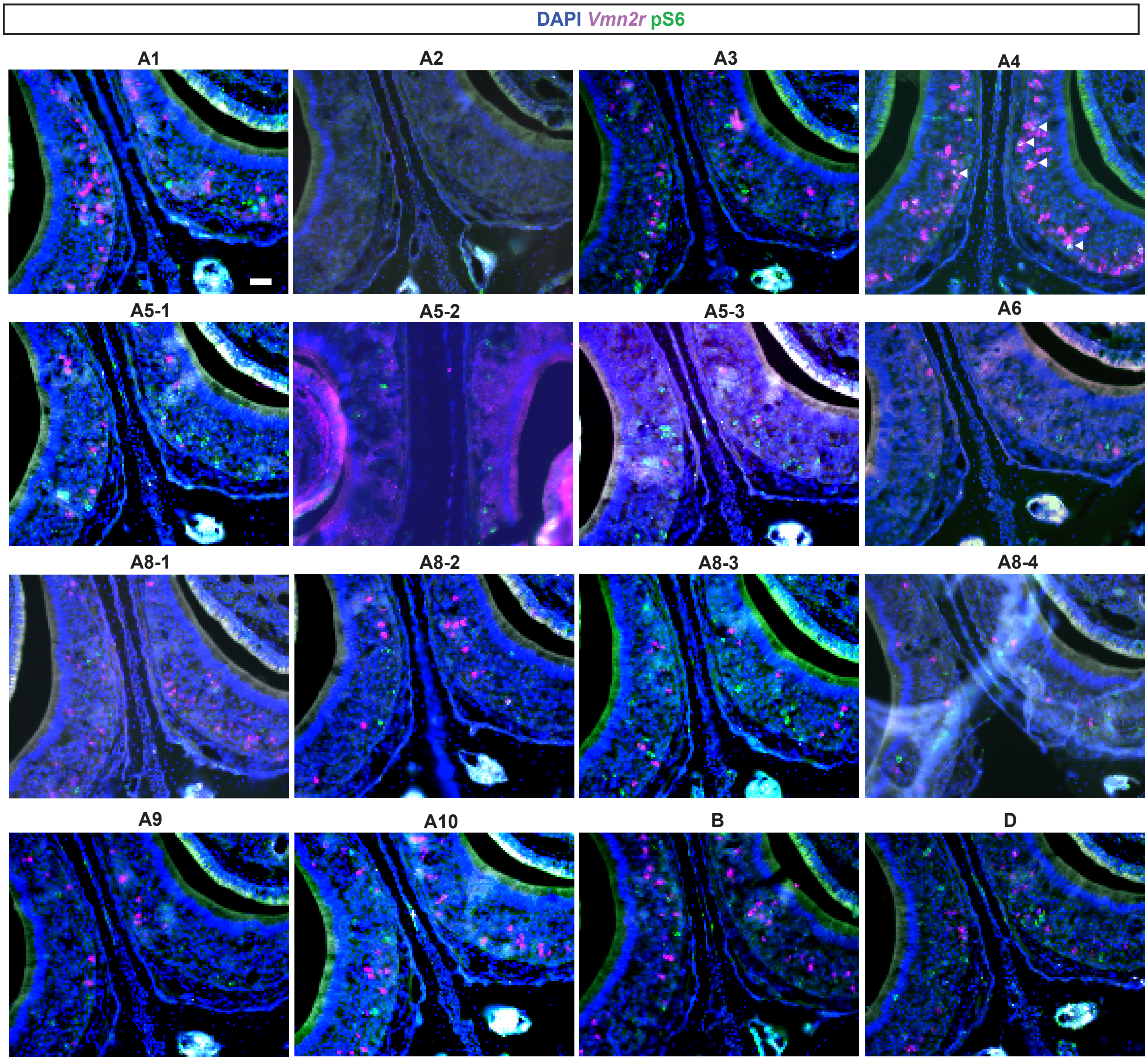


**Supplemental Figure 3**

Representative images of V2R in situ hybridization and pS6 immunohistochemistry for each V2R subfamily.
