## Supplemental Figure 4 for "Hypothalamic representation of the imminence of predator threat detected by the vomeronasal organ in mice"

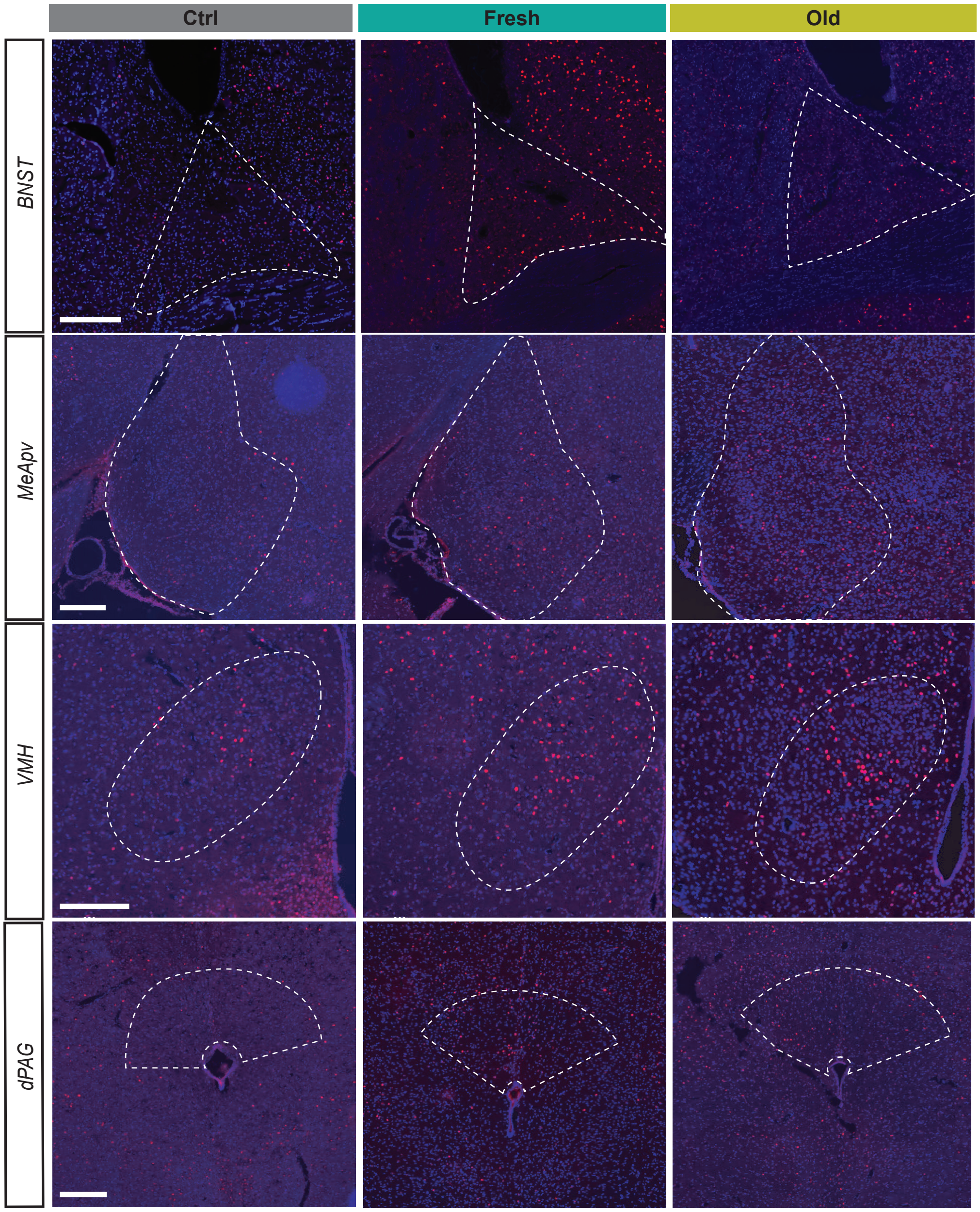


**Supplemental Figure 4**

Representative images displaying the expression of cFos (red) in the BNST, MeApv, VMH, and dPAG of mice exposed to control, fresh, or old saliva swabs. Each neural substrate is delineated by a dashed white line, defined by counterstaining with DAPI (blue). Scale bar: 200 µm.
