## Supplemental Figure 5 for "Hypothalamic representation of the imminence of predator threat detected by the vomeronasal organ in mice"

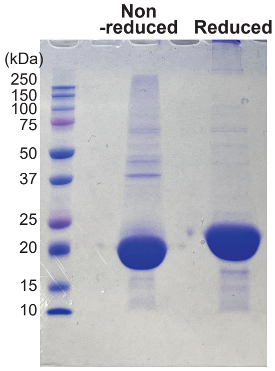


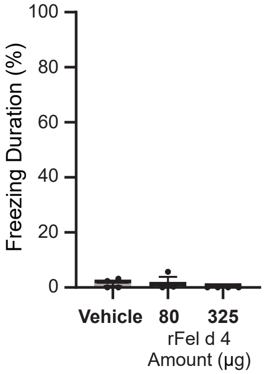


**Supplemental Figure 5**

(Left) SDS-PAGE gel image showing 65 µg of recombinant Fel d 4 (rFel d 4) protein in non-reduced and reduced conditions. (Right) Percentage of total freezing episodes directed toward swabs containing vehicle, 80 µg Fel d 4, or 325 µg Fel d 4. Sample sizes are as follows: Vehicle (n = 4), 80 µg Fel d 4 (n = 3), and 325 µg Fel d 4 (n = 4).
