## Supplemental Table 1 for "Hypothalamic representation of the imminence of predator threat detected by the vomeronasal organ in mice"

A

| Freezing (Figure 1K) | |
|---|---|
| | Adjusted P Value |
| Fresh vs. Old | 0.0002 |
| Fresh vs. 70% Fresh | 0.9638 |
| Fresh vs. 50% Fresh | 0.0014 |
| Fresh vs. 30% Fresh | 0.018 |
| Old vs. 70% Fresh | 0.0063 |
| Old vs. 50% Fresh | >0.9999 |
| Old vs. 30% Fresh | 0.9694 |
| 70% CS vs. 50% Fresh | 0.0104 |
| 70% CS vs. 30% Fresh | 0.0466 |
| 50% CS vs. 30% Fresh | 0.9681 |

B

| Two-way ANOVA | | | |
|---|---|---|---|
| Section | Variation | F (DFn, DFd) | P value |
| 1 | bin x swab type | F (18, 162) = 3.930 | <0.0001 |
| | bin | F (1.795, 32.30) = 13.63 | <0.0001 |
| | swab type | F (2, 18) = 5.753 | 0.0117 |
| 2 | bin x swab type | F (18, 234) = 2.538 | 0.0008 |
| | bin | F (2.961, 76.99) = 20.84 | <0.0001 |
| | swab type | F (2, 26) = 7.364 | 0.0029 |
| 3 | bin x swab type | F (18, 288) = 8.220 | <0.0001 |
| | bin | F (2.684, 85.89) = 32.86 | <0.0001 |
| | swab type | F (2, 32) = 14.75 | <0.0001 |
| 4 | bin x swab type | F (18, 252) = 3.347 | <0.0001 |
| | bin | F (1.595, 44.67) = 16.10 | <0.0001 |
| | swab type | F (2, 28) = 5.491 | 0.0097 |
| 5 | bin x swab type | F (18, 234) = 2.937 | <0.0001 |
| | bin | F (1.964, 51.06) = 12.23 | <0.0001 |
| | swab type | F (2, 26) = 4.325 | 0.0239 |

| *p* value (Tukey's multiple comparisons test) | | | | | | |
|---|---|---|---|---|---|---|
| Bin | | Section 1 | Section 2 | Section 3 | Section 4 | Section5 |
| 1 | Ctrl vs. Fresh | 0.052 | 0.9767 | 0.0324 | 0.0269 | 0.0367 |
| | Ctrl vs. Old | 0.4929 | 0.7034 | 0.0316 | 0.1803 | 0.0281 |
| | Fresh vs. Old | 0.6752 | 0.584 | 0.9954 | 0.8789 | 0.6207 |
| 2 | Ctrl vs. Fresh | 0.1021 | 0.2383 | 0.0013 | 0.0396 | 0.0598 |
| | Ctrl vs. Old | 0.3855 | 0.2196 | 0.115 | 0.0527 | 0.1092 |
| | Fresh vs. Old | 0.9042 | 0.7204 | 0.2985 | 0.359 | 0.5757 |
| 3 | Ctrl vs. Fresh | 0.1046 | 0.0187 | <0.0001 | 0.0225 | 0.0286 |
| | Ctrl vs. Old | 0.0764 | 0.0144 | 0.0726 | 0.1169 | 0.0172 |
| | Fresh vs. Old | 0.9102 | 0.9756 | 0.0064 | 0.8327 | 0.7651 |
| 4 | Ctrl vs. Fresh | 0.0916 | 0.0021 | 0.0005 | 0.0504 | 0.0672 |
| | Ctrl vs. Old | 0.179 | 0.0523 | 0.129 | 0.0577 | 0.0146 |
| | Fresh vs. Old | 0.6076 | 0.9851 | 0.0141 | 0.3502 | 0.9129 |
| 5 | Ctrl vs. Fresh | 0.0539 | 0.0067 | 0.0715 | 0.1121 | 0.2498 |
| | Ctrl vs. Old | 0.363 | 0.9797 | 0.7531 | 0.0514 | 0.1533 |
| | Fresh vs. Old | 0.2223 | 0.0035 | 0.4166 | 0.9068 | 0.7722 |
| 6 | Ctrl vs. Fresh | 0.0214 | 0.4539 | 0.5051 | 0.0568 | 0.0969 |
| | Ctrl vs. Old | 0.4455 | 0.7104 | 0.2474 | 0.2551 | 0.1941 |
| | Fresh vs. Old | 0.0849 | 0.1688 | 0.9157 | 0.911 | 0.5344 |
| 7 | Ctrl vs. Fresh | 0.035 | 0.003 | 0.0525 | 0.6027 | 0.0848 |
| | Ctrl vs. Old | 0.6353 | 0.1991 | 0.1411 | 0.805 | 0.3781 |
| | Fresh vs. Old | 0.2889 | 0.5182 | 0.9863 | 0.8601 | 0.9728 |
| 8 | Ctrl vs. Fresh | 0.119 | 0.0284 | 0.0268 | 0.4311 | 0.7418 |
| | Ctrl vs. Old | 0.427 | 0.2746 | 0.3347 | 0.269 | 0.7179 |
| | Fresh vs. Old | 0.8121 | 0.8054 | 0.9744 | 0.9737 | 0.9905 |
| 9 | Ctrl vs. Fresh | 0.799 | 0.1698 | 0.1948 | 0.3013 | 0.5455 |
| | Ctrl vs. Old | 0.467 | 0.6465 | 0.8618 | 0.8703 | 0.6078 |
| | Fresh vs. Old | 0.5811 | 0.8893 | 0.4167 | 0.2078 | 0.9382 |
| 10 | Ctrl vs. Fresh | 0.3358 | 0.0593 | 0.39 | 0.5308 | 0.439 |
| | Ctrl vs. Old | 0.3892 | 0.1434 | 0.3244 | 0.3397 | 0.9736 |
| | Fresh vs. Old | 0.1655 | 0.979 | 0.7183 | 0.9532 | 0.3543 |

**Supplemental Table 1**

(**A**) Statistical summary of Tukey's multiple comparisons for Figure 1K. (**B**) Statistical summary of the two-way ANOVA with Tukey's multiple comparisons for Figure 6B.
